# No evidence for insecticide resistance in a homogenous population of *Aedes albopictus* in Mecklenburg County, North Carolina

**DOI:** 10.1101/2020.06.05.136135

**Authors:** Stephanie J. Mundis, Gabriela Hamerlinck, Emily K. Stone, Ari Whiteman, Eric Delmelle, Tyler Rapp, Michael Dulin, Sadie J. Ryan

## Abstract

*Aedes albopictus* is a cosmopolitan mosquito species capable of transmitting arboviral diseases such as dengue, chikungunya, and Zika. To control this and similar species, public and private entities often rely on pyrethroid insecticides. Insecticide resistance status and physiological traits, such as body size, may contribute to local patterns of abundance, which is important for planning vector control. In this study, we genetically screened *Ae. albopictus* collected from June to August, 2017, in Mecklenburg County, North Carolina, for mutations conferring pyrethroid resistance, and examined spatiotemporal patterns of specimen size, as measured by wing length. We hypothesized that size variation would be associated with factors found to influence abundance in similar populations of *Ae. albopictus*, and could therefore serve as a proxy measure. The genetic screening results indicated that known pyrethroid resistance alleles in two *kdr* regions are not present in this population. We detected no significant associations between wing length and socioeconomic and landscape factors, but mosquitoes collected in June had significantly longer wing length than in July or August. The lack of resistance indicators suggest that this population has not developed insecticide resistance via voltage-gated sodium channel mutations. The greater wing lengths in June are likely driven by meteorological patterns, suggesting that short-term weather cues may modulate morphological characteristics that, in turn, affect local fecundity and virus transmission potential.

## Introduction

Recent emergences and spread of diseases such as dengue, chikungunya, and Zika, have led to an uptick in public interest and concern about vector control for public health in the United States. Greater knowledge about the presence and distribution of disease vectoring *Aedes ssp*. mosquitoes can inform and guide vector control. However, an additional factor has emerged in recent years: insecticide resistance. This study undertook an examination of vector surveillance data for *Aedes albopictus* mosquitoes in Mecklenburg County, North Carolina to assess potential factors affecting distribution, and evidence for the emergence of insecticide resistance.

First identified in Texas in 1985, the invasive *Ae. albopictus* has dramatically expanded its range in the United States (1,2). This U.S. expansion is part of a larger global trend: in the last 50 years, *Ae. albopictus* has spread to all inhabited continents (3), and has become established in both tropical and temperate environments (1). This species is a container-breeder and feeds opportunistically, biting a wide range of hosts, although some populations exhibit a preference for mammals, and, more specifically, humans (4). While *Ae. albopictus* is less anthropophilic than *Ae. aegypti*, it can serve as a vector for the same diseases as *Ae. aegypti*, including dengue, chikungunya, Rift Valley fever, yellow fever, and Zika (5). Moreover, the opportunistic feeding behavior of *Ae. albopictus* may allow this species to act as a bridge vector, facilitating spillover of zoonotic diseases into human populations (4). Additionally, populations of *Ae. albopictus* have been found to competitively displace populations of *Ae. aegypti* (6).

In light of the disease transmission role of *Ae. albopictus*, its recent global expansion, and its ability to out-compete other important vector species, it is unsurprising that this species is of concern to public health and been accordingly targeted by vector control programs. *Ae. albopictus* was an important vector in the 2004-2007 chikungunya epidemic across several Indian Ocean islands (7), and the 2007 concurrent outbreak of dengue and chikungunya in Gabon (8) and additional chikungunya outbreaks in Italy in 2007 (9) and 2017 (10). *Ae. albopictus* has also been implicated as a vector of La Crosse virus (11), the most common endemic arboviral disease in North Carolina, where this study takes place (12). *Aedes albopictus* control often relies on the use of adulticides (13), as is the case in North Carolina, where pyrethroid insecticides are commonly used for barrier spraying to control *Ae. albopictus* (14). While insecticide resistance has been documented in *Ae. aegypti* populations around the world, fewer studies have focused on the resistance status of *Ae. albopictus*, with most work on this species concentrated in Southeast Asia, where resistance to all four major insecticide classes has been reported (5). In the United States, however, *Ae. albopictus* populations remain broadly susceptible to most insecticide treatments, though low levels of resistance to organophosphates and dichlorodiphenyltrichloroethane (DDT) have been detected in Florida and New Jersey populations (15).

*Ae. albopictus* abundance can vary across space and time, influencing local disease transmission potential. Socioeconomic, landscape, and seasonal factors have been associated with *Ae. albopictus* abundance; areas of low socioeconomic status (SES) often have a greater number of discarded containers available for *Ae. albopictus* breeding (16) and pupae from *Aedes* species are more likely to be found in neighborhoods below median income (17). This was demonstrated recently in Mecklenburg County, North Carolina, in which abundance of gravid *Ae. albopictus* was found to be significantly higher in low-income neighborhoods (18). This work identified land cover factors associated with increased *Ae. albopictus* abundance, including the percent of land covered by buildings, tree canopy, grass and shrubs, roads and railroads, and the overall diversity of land cover types in a 30-meter buffered area around sampled sites (18, 19). Previous studies found that small patches of vegetation in urban areas, such as parks, gardens, and playgrounds, are often associated with high *Ae. albopictus* abundance (20), and that peaks in abundance often occur in late summer months in temperate climates (18,20,21).

This study aimed to establish a baseline description of the insecticide resistance status and patterns of morphological variation within a population of *Ae. albopictus* collected from Mecklenburg County, North Carolina. As such, our first objective was to screen adult *Ae. albopictus* females for genetic mutations indicating resistance to pyrethroids, the most commonly used class of insecticides for barrier spraying in North Carolina (14). Our second objective was to assess whether a morphological measure could act as a proxy for abundance, reducing the need and cost for time-consuming direct abundance sampling. Wing size is correlated to body size in *Ae. albopictus*, and we hypothesized that variation in female *Ae. albopictus* size is associated with the socioeconomic, landscape, and seasonal factors that influence *Ae. albopictus* abundance. Since multiple studies have found that female *Ae. albopictus* size is positively correlated with fecundity (22,23), we predicted we would observe larger mean wing lengths in the socioeconomic classes, land cover types, and time periods associated with higher *Ae. albopictus* abundance. Furthermore, vector size influences virus transmission potential, with viral dissemination more likely in smaller individuals, as has been shown for dengue virus in *Ae. albopictus* (24) and La Crosse virus in *Aedes triseriatus* (25). If size can serve as a reliable proxy for either abundance or disease transmission potential in a local context, it provides a low-cost means to prioritize and target areas of importance, rather than time-consuming abundance sampling measures.

## Materials and Methods

### Study site

We performed our analyses using adult female *Ae. albopictus* collected from June to August 2017 in Mecklenburg County, North Carolina, which encompasses the city of Charlotte (Figure 1). Mecklenburg County had an average population density of approximately 1,900 people per square mile, and a median household income of $61,695, with 13.4% of the population classified as persons in poverty in 2017 (26). Of the 50 largest cities in the United States, Charlotte had the lowest rate of upward economic mobility in the 2000 census (27) and has also been characterized by pervasive racial segregation and high rates of income inequality (18,28). The *Ae. albopictus* specimens used in this study were collected from 90 unique sampling sites selected to maximize spatial distribution across the county and to represent the range of values present across a variety of socioeconomic and landscape factors, as described in Whiteman et al. (18).

**Figure 1:**
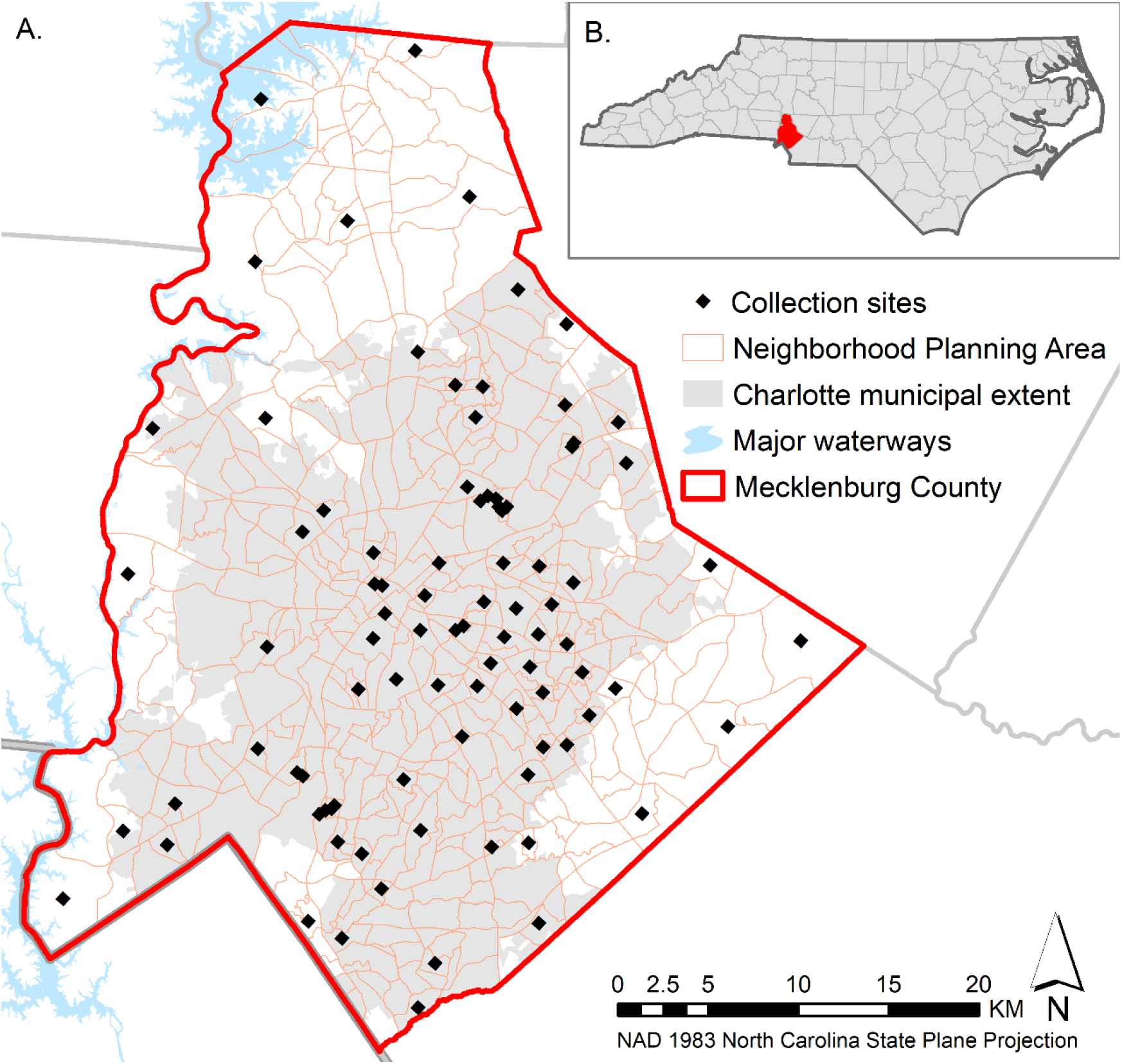
A. Collection sites in Mecklenburg County, North Carolina. B. Location of Mecklenburg County in North Carolina.

### DNA extraction, amplification, and sequencing

We destructively extracted DNA from whole mosquitoes for use in PCR using Qiagen DNeasy isolation kits (Qiagen Sciences, Germantown, MD, USA). For all samples, we amplified and sequenced two regions of *kdr* (domain II, 381 bp; domain IV, 280 bp) using the AegSCF20/AegSCR21 and AlbSCF6/AlbSCR8 primer pairs, respectively (Supplemental Table 1). Amplification of *kdr* domain III was unsuccessful, despite multiple attempts and consultations with other researchers conducting resistance screening in *Aedes spp*. The thermocycler conditions were identical for *kdr* domains II and IV, an initial denaturing step at 96°C for 10 min, 40 cycles of 30 s at 96°C, 30 s at 55°C and 45 s at 72°C; with a final extension step of 10 min at 73°C (Supplemental Table 1). All mosquito PCR products were cleaned using exonuclease I and shrimp alkaline phosphatase (Fisher Scientific, Pittsburgh, PA). Sequencing was performed by GeneWiz. We used MegaX (29) and BioEdit (30) to assemble and form contigs of our forward and reverse reads.

### Wing length measurements

We aimed to measure the wing length of one *Ae. albopictus* adult female from each of the 90 collection sites for each month in the collection window. However, because some sites did not yield *Ae. albopictus* females each month, or specimens were in poor condition, we measured 236 wings total (representing 84, 72, and 80 sites in June, July, and August, respectively). We used a camera mounted on a dissecting microscope to photograph mosquito wings. We then processed all images with ImageJ (31) to measure the length of each wing. Each wing was measured by two authors (SM and ES) independently, and measurements averaged to determine a consensus length. If there was a difference greater than 2 mm between the independent measurements, a third measurement was taken by a third author (GH) and computed into the average for the month.

To test for statistically significant associations between socioeconomic variables and wing length, we first tested for wing length differences across socioeconomic quintiles based on the 2016 median household income at the Neighborhood Planning Area (NPA) level. The NPA is a unit developed by the Charlotte-Mecklenburg Planning Commission which approximates the census tract, but with improved representation of actual neighborhoods within the county (18). Mean wing length measurements across the sampling period per site were tested for normality using the Shapiro-Wilk test, and were found not to be normally distributed (p value < 0.004). We conducted a Kruskal-Wallis test and pairwise Wilcoxon rank sum test to assess differences in wing length across income groups. We tested for associations between mean wing length and socioeconomic or human demographic variables at the NPA level previously found to be related to *Ae. albopictus* abundance (18), using Spearman rank correlations. These variables included: violent crime rate, population density, employment rate, proportion Hispanic population, foreclosure rate, proximity to a park, and proportion African-American population (18).

We used the 2012 Mecklenburg County Tree Canopy/Land Cover dataset to test for associations between land cover and wing length. This dataset was developed at a 3.33-foot spatial resolution using object-based image analysis techniques along with 2012 LiDAR data, 2012 National Agriculture Imagery Program imagery, and ancillary spatial datasets (32). Land cover types included buildings, roads/railroads, tree canopy, grass/shrubs, water, and other paved surfaces. We generated a 30-meter buffer around each sampling site and calculated the percentage of each land cover type present within each buffer. Previous work has indicated that a 30-meter buffer is the best scale to detect the relationship between high-resolution land cover variables and *Aedes* abundance (33). We tested for correlations between percent of each land cover type present within the buffer and mean wing length at each collection site using Spearman rank correlations. Additionally, we used a Kruskal-Wallis test to identify significant differences in mean wing length at sites classified as rural (n=5), suburban (n=63), and urban (n=20) for each collection month and the total sampling season. These designations were based on percent impervious surface (roads/railroads, other paved surfaces, and buildings) within the 30-meter buffer, using cut-off values from Murdock et al (34). We tested for statistically significant differences in mean wing length across collection months using a Kruskal-Wallis test, and post hoc Wilcoxon rank sum tests for pairwise differences. We used Global Moran’s I tests to detect spatial autocorrelation in the mean wing lengths across the study area for each month and for the averaged wing lengths for the entire sampling period, using inverse distance to conceptualize spatial relationships. All statistical tests were performed in R v3.5.0 (35), and all spatial data processing was conducted in ArcGIS 10.6 (36).

## Results

We successfully extracted DNA from 86 mosquitoes, representing 95% of the total 90 collection sites. Amplification and sequencing of *kdr* domains II and IV were successful for 27 individuals (30% coverage) and 75 individuals (83% coverage), respectively. Amplification of *kdr* domain III was unsuccessful. We found no mutations that would infer pyrethroid resistance among our samples from Mecklenburg County, NC. The resulting sequences for all samples were deposited in GenBank; accession numbers can be found in Supplemental Table 2.

The mean wing length for the 236 female *Ae. albopictus* specimens was 2.73 mm (range 1.64 mm to 4.29 mm). Differences in wing length across median income quintiles were not statistically significant (Kruskal-Wallis χ^2^=2.645, df=4, p=0.6189). The Spearman rank correlations between wing length and the socioeconomic and human demographic variables previously identified as being associated with *Ae. albopictus* abundance (18), were not significant.

Wing length was not significantly associated with land cover factors. This included the Spearman’s rank correlation between the percentage of each land cover type present within the 30-meter buffer around each sampling site. There was also no significant difference in mean wing length across rural, suburban, and urban areas, based on percent impervious surface (Kruskal-Wallis χ^2^=1.2745, df=2, p-value=0.5287).

Differences in mean wing length across the three sampling months was statistically significant (Figure 2; Kruskal-Wallis χ^2^=9.9499, df=2, p-value=0.0069). We found a statistically significant difference between wing length measurements of samples collected in June and August (Wilcoxon rank sum test, p-value=0.008) and between June and July (Wilcoxon rank sum test, p-value = 0.022), but not August and July (Wilcoxon rank sum test, p-value=0.54). The mean wing lengths of collected mosquitoes was longest in June, indicating the June collections had the largest females. The Global Moran’s I tests for spatial autocorrelation did not indicate statistically significant clustering or dispersion in mean wing lengths when averaged over the entire study period (p-value = 0.67), not for each month individually (June p-value = 0.98; July p-value = 0.27; August p-value = 0.73).

**Figure 2:**
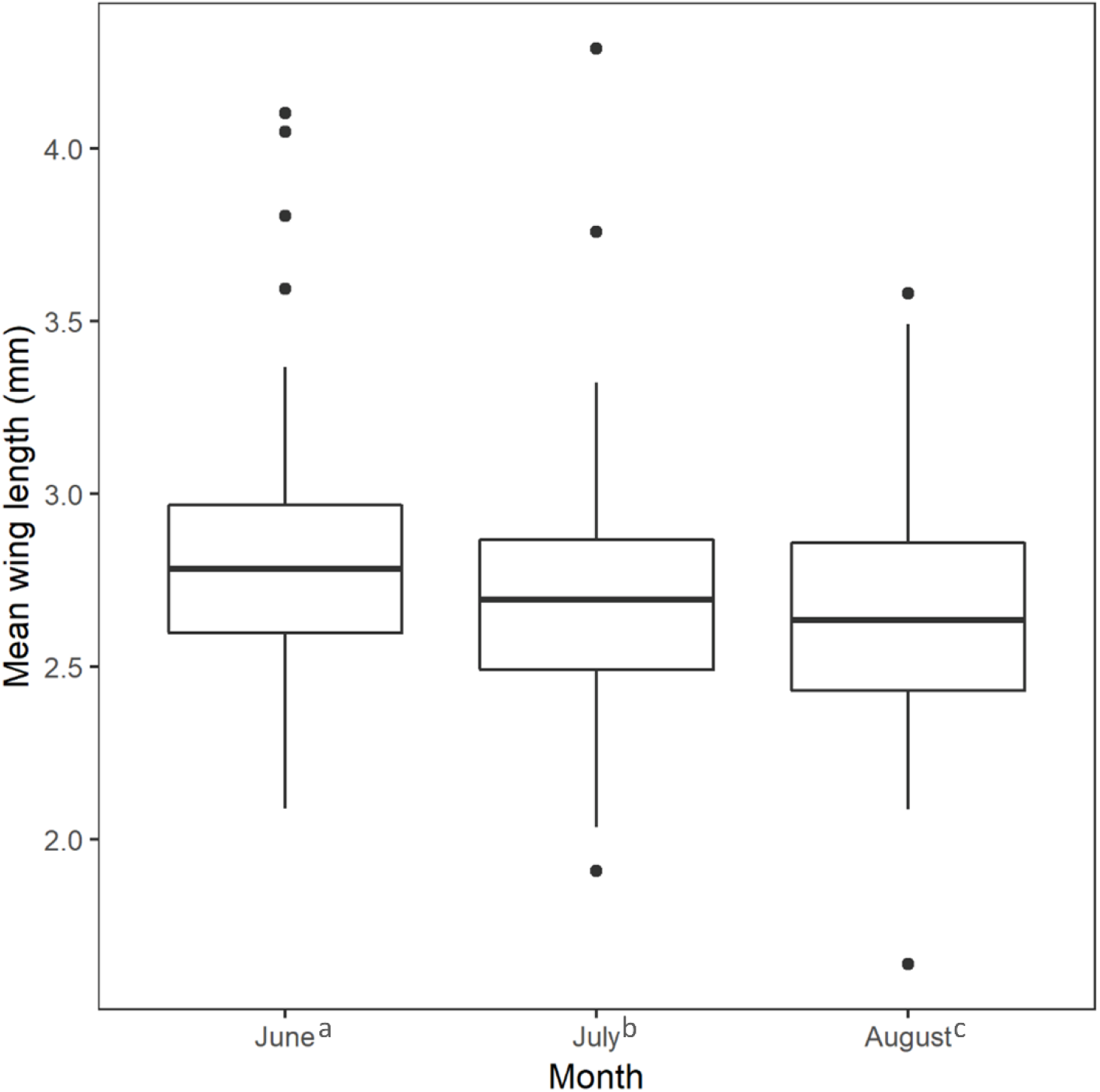
Mean female *Aedes albopictus* wing length by collection month. The mean June wing length was longer than July and August mean wing lengths. There was no difference between July and August mean wing length. Superscript lowercase letters indicate values were significantly different from one another in Kruskal-Wallis tests with a post hoc Wilcoxon rank sum test at p ≤ 0.05.

## Discussion

As the range of *Ae. albopictus* continues to expand, continuous surveillance and study of the species is needed. Regular monitoring of insecticide susceptibility is essential to promptly identify the emergence of resistance and implement appropriate, alternative control measures (37). Having a baseline understanding of the morphology and distribution of vector populations within the context of local socioeconomic, landscape, and temporal influences can inform targeted abatement strategies. In this study, we screened *Ae. albopictus* collected from Mecklenburg County, North Carolina, for genetic indicators of resistance, and examined spatial and temporal patterns of wing length variation among the collected adult female *Ae. albopictus* specimens.

While our sample size was relatively small, the homogenous lack of genetic indicators of pyrethroid resistance across the study area suggests that this population is broadly susceptible to pyrethroid insecticides, consistent with findings from similar studies in this area. In a 2018 study, researchers found that *Ae. albopictus* populations from seven North Carolina counties, including Mecklenburg County, were susceptible to five commonly used pyrethoids in CDC bottle bioassays, with the exception of Pitt County, where developing resistance (93% mortality) to permethrin was documented (14). In contrast, resistance to chlorpyrifos and malathion, two commonly used organophosphates, was documented in all seven populations in the same study (14). Budgets for mosquito control programs in North Carolina have been dramatically reduced in the past decade (38) and a recent survey found that approximately 31% of respondents in North Carolina personally administer insecticides for mosquito control on their property (39). This combination of limited resources for oversight and unregulated insecticide applications by private individuals indicates that selection for insecticide resistant mosquitoes will likely continue in this area, although more information is needed to predict whether pyrethroid resistance will develop.

We did not detect significant associations between *Ae. albopictus* wing length and most of the socioeconomic and landscape factors considered in this study, although these factors were associated with *Ae. albopictus* abundance in other studies. This means that our hypothesis that larger female *Ae. albopictus* females with higher fecundity drive increases in local abundance was not supported. While certain areas may produce larger female *Ae. albopictus* that have more offspring, their impact on local abundance is likely hampered by high larval densities that result in smaller adults (40). Furthermore, the Global Moran’s I tests indicate that wing length had a random distribution, indicating that female adult size is likely the result of multiple random, interacting processes. We did not observe significant differences in wing length between rural, suburban, and urban sites. This is contradictory to recent work conducted in Athens, Georgia (34), which showed that *Ae. albopictus* emerging from containers placed in urban sites were significantly smaller than those emerging in rural sites (34). However, this difference was statistically significant only in the fall, and our study was limited to a single summer season of collections. Differences in wing length across land cover types might be found if sampling in Mecklenburg County were to continue into the fall.

We found that *Ae. albopictus* collected in June had significantly longer wing lengths than *Ae. albopictus* collected in August and July, which could be due to meteorological conditions. The total monthly precipitation for Charlotte in June, July, and August, 2017 was 4.3 inches, 4.45 inches, and 5.29 inches respectively (41). Observing significantly larger mosquitoes during the month with the lowest rainfall corresponds to a 2001 study showing that *Ae. albopictus* reached their largest size in an environment where their water source was allowed to evaporate completely (42). The temperature was lower in June than in July or August during the study period, with an average high of 29.8°C in June, compared to 33.1°C in July, and 30.8°C in August. Perhaps counter-intuitively, higher temperatures have been shown to result in the growth of heavier adult *Ae. albopictus* with shorter wings (43); given this temperature dependent growth is likely curvilinear, it is also plausible that it is pushed beyond optimal growth temperatures in July and August. Further work is needed to explore how these meteorological variables interact with each other and other environmental factors to determine *Ae. albopictus* size.

In conclusion, this work served to establish a baseline description for the *Ae. albopictus* population in Mecklenburg County, North Carolina. We did not detect indicators of insecticide resistance, but continued surveillance remains critical for early detection of diminished susceptibility to insecticide. Similarly, further research will likely illuminate the extent to which spatial and temporal factors influence variation in wing size in this and other similar populations, and the utility of wing size measures for application outside the laboratory. At this scale of collection, we were unable to establish a statistically reliable indicator of variation in abundance across space, using a morphological measurement.

## Acknowledgements

The contributions of SJR, SJM, and GH were supported by CDC grant 1U01CK000510-01: Southeastern Regional Center of Excellence in Vector-Borne Diseases: the Gateway Program. This publication was supported by the Cooperative Agreement Number above from the Centers for Disease Control and Prevention. Its contents are solely the responsibility of the authors and do not necessarily represent the official views of the Centers for Disease Control and Prevention. ES was supported by the UF Emerging Scholars program. Fieldwork was funded by the Academy for Population Health Innovation, a collaboration between Mecklenburg County Public Health and The University of North Carolina at Charlotte.

## Tables

**Supplemental Table 1:**
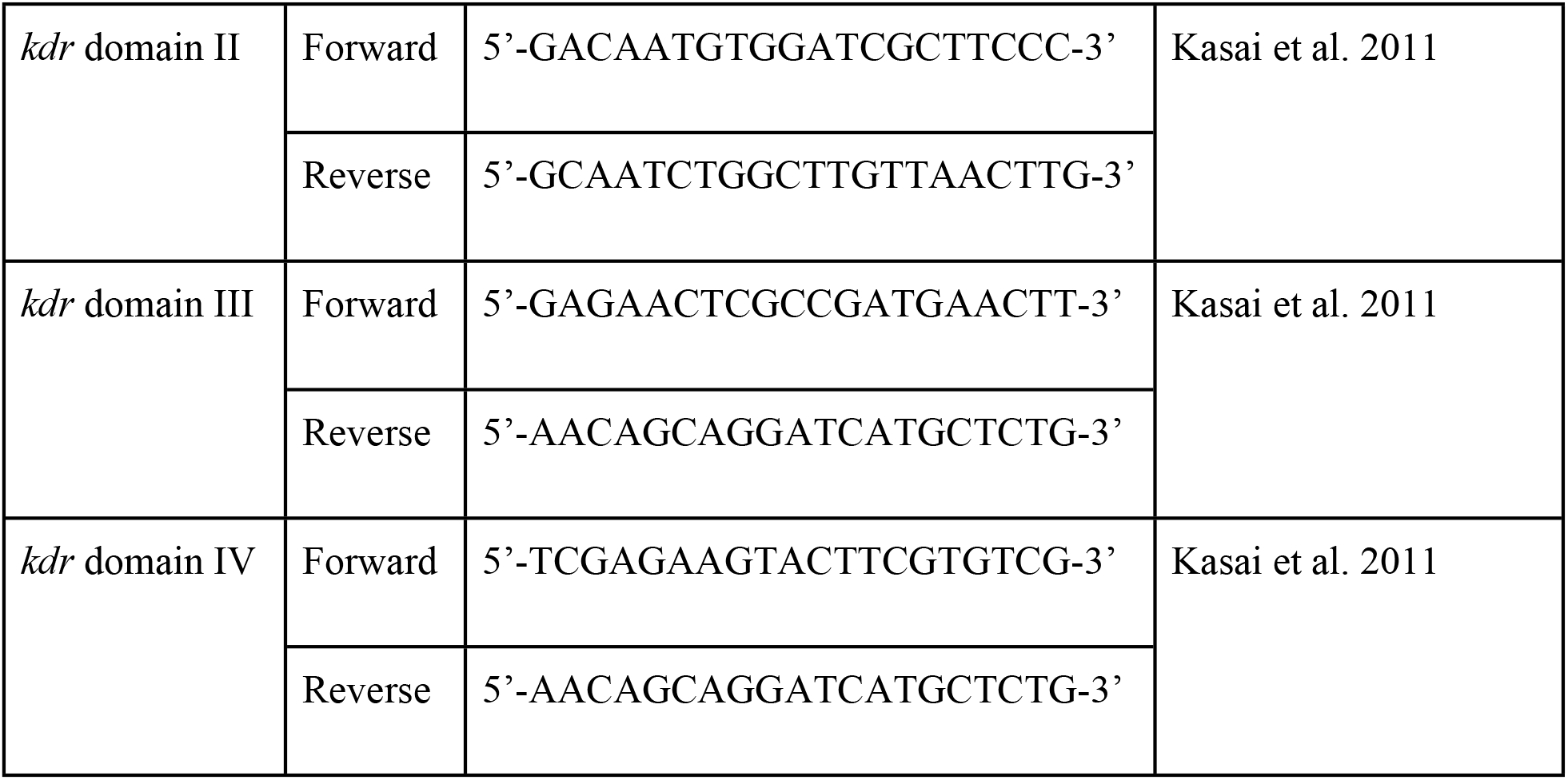
Primers used for amplification of three domains of *kdr*.

**Supplemental Table 2:** Accession numbers for *kdr* domains II and IV.

| Site ID | Neighborhood<br>classification | *kdr* domain II | *kdr* domain IV |
| --- | --- | --- | --- |
|  |  | Accession ## | Accession ## |

